# Three-dimensional electro-neural interfaces electroplated on subretinal prostheses

**DOI:** 10.1101/2023.11.09.566003

**Authors:** Emma Butt, Bing-Yi Wang, Andrew Shin, Zhijie Charles Chen, Mohajeet Bhuckory, Sarthak Shah, Ludwig Galambos, Theodore Kamins, Daniel Palanker, Keith Mathieson

## Abstract

**Objective:** High-resolution retinal prosthetics offer partial restoration of sight to patients blinded by retinal degenerative diseases through electrical stimulation of the remaining neurons. Decreasing the pixel size enables an increase in prosthetic visual acuity, as demonstrated in animal models of retinal degeneration. However, scaling down the size of planar pixels is limited by the reduced penetration depth of the electric field in tissue. We investigate 3-dimensional structures on top of the photovoltaic arrays for enhanced penetration of electric field to permit higher-resolution implants.

**Approach:** We developed 3D COMSOL models of subretinal photovoltaic arrays that accurately quantify the device electrodynamics during stimulation and verified it experimentally through comparison with the standard (flat) photovoltaic arrays. The models were then applied to optimise the design of 3D electrode structures (pillars and honeycombs) to efficiently stimulate the inner retinal neurons. The return electrodes elevated on top of the honeycomb walls surrounding each pixel orient the electric field inside the cavities vertically, aligning it with bipolar cells for optimal stimulation. Alternatively, pillars elevate the active electrode into the inner nuclear layer, improving proximity to the target neurons. Modelling results informed a microfabrication process of electroplating the 3D electrode structures on top of the existing flat subretinal prosthesis.

**Main results:** Simulations demonstrate that despite the conductive sidewalls of the 3D electrodes being exposed to electrolyte, most of the charge flows via the high-capacitance sputtered Iridium Oxide film that caps the top of the 3D structures. The 24 µm height of the electroplated honeycomb structures was optimised for integration with the inner nuclear layer cells in rat retina, while 35 µm height of the pillars was optimized for penetrating the debris layer in human patients. Release from the wafer and implantation of the 3D arrays demonstrated that they are mechanically robust to withstand the associated forces. Histology demonstrated successful integration of the 3D structures with the rat retina in-vivo.

**Significance:** Electroplated 3D honeycomb structures produce a vertically oriented electric field that offers low stimulation threshold, high spatial resolution and high contrast for the retinal implants with pixel sizes down to 20µm in width. Pillar electrodes offer an alternative configuration for extending the stimulation past the debris layers. Electroplating of the 3D structures is compatible with the fabrication process of the flat photovoltaic arrays, thereby enabling much more efficient stimulation than in their original flat configuration.

## 1. Introduction

Age-related macular degeneration (AMD) is one of the leading causes of irreversible sight loss worldwide [1], affecting an estimated 200 million patients. In its atrophic form, called geographic atropy (GA), this degenerative retinal condition leads to loss of the photoreceptor cells in the central macula [2], the high resolution region of the retina responsible for our central vision, thus impairing patients’ ability to read and recognize faces. Despite the loss of photoreceptors, the inner retinal neurons can remain functional, and electrical stimulation of these neurons can evoke visual percepts [3]. Recent clinical trials with a subretinal photovoltaic array PRIMA (Pixium Vision, Paris, France) demonstrated form perception in GA of AMD patients, with prosthetic acuity reaching the level of 20/438, closely matching the implant’s pixel size of 100 µm, which corresponds to the acuity limit of 20/420 [4]. Since the remaining peripheral vision in AMD patients often supports visual acuity of no worse than 20/400, clinically meaningful improvement requires smaller pixels. For example, a visual acuity exeeding 20/100 would require pixels of about 20 µm [5]. Patterned electrical stimulation of the retina with 20 µm pixels has demonstrated a grating acuity up to the natural resolution limit of 27 µm in rats [25]. However, new strategies are needed to safely translate this to a significantly thicker human retina [20]. Subretinal implants aim to activate the bipolar cells in the inner nuclear layer [3] by polarizing them in electric field, and then rely on the remaining retinal neural network to process their output and evoke the bursts of action potentials in the retinal ganglion cells. Utilizing this remaining retinal network has been shown to preserve many features of the retinal signal processing, including flicker fusion, antagonistic center-surround, and others [6].

In the PRIMA system, the near-IR pulses (880 nm) projected onto the photovoltaic implant from the augmented-reality glasses are converted into pulses of electric current, injected into electrolyte via the active electrodes in each pixel and collected by the return electrodes surrounding each pixel. Decreasing the pixel size can increase the achievable visual acuity, but stimulation thresholds rapidly increase [7] due to reduced penetration of E-field into the tissue and reduced photosensitive area in each pixel. They can be compensated by higher IR irradiance, but for pixels smaller than 40 µm in rodents and 75 µm in humans, the required irradiance exceeds the ocular safety limit for near-IR exposure (8.25 mW/mm^2^ at 10 ms pulse duration and 30 Hz repetition rate) [8]. Stronger stimuli are required with human retina because it is thicker than in rodents and because it exhibits a 35 µm subretinal debris layer in atrophic areas, which increases the separation between the target cells and the implant [9].

3D electrode structures offer a solution to this problem, as the stimulating electric field can either be shaped for more efficient stimulation or brought closer to the target neurons. Previous studies with passive 3D implants demonstrated that inner retinal neurons migrate into the voids in the implant, and thereby can achieve close proximity to electrodes [10-13]. Two types of 3D electrode structures have been proposed: a raised return electrode in a hexagonal array (so-called honeycombs) [11] and pillar electrodes that raise the active electrode to the target neuronal layer [10]. Both approaches have advantages and limitations. For example, the honeycomb structures align the electric field vertically within the well, matching the dominant orientation of bipolar cells, thus reducing their stimulation threshold and decreasing the pixel-to-pixel cross-talk. However, it is unclear how such structures will integrate with a debris layer in human retina. Pillar electrodes, on the other hand, may penetrate through this debris layer, bringing the stimulation site close to the target inner retinal neurons. However, the spread of current from the pillar top is more spherical, so that the threshold and contrast may be degraded, compared to honeycombs. Previously, we investigated short (10 µm) pillars in RCS rats, where there is no subretinal debris, and observed a moderate (2-fold) reduction in stimulation threshold with 55µm pixels [14]. The pixels investigated here are much smaller – down to 20µm, and pillars are much taller (35µm in height), designed to raise the active electrode above the debris layer between implant and the INL in humans [15], and thus a much more significant reduction of the stimulation threshold is expected.

These high-aspect ratio electrode structures present a microfabrication challenge, and we describe electroplating process for such 3D electrodes on a photovoltaic implant. The structures are modelled using 3D finite element analysis (COMSOL Multiphysics 5.6 with electrochemistry and circuit modules). This model was first verified by comparison with experimental results from a Pixium PRIMA chip (Fig. 1D). It was then extended to model an array of conductive 3D structures acting as return electrodes on honeycombs or active electrodes on pillars. This model informed the fabrication process for both of these devices, highlighting the effect of the low-capacitance side walls and the high-capacitance top coating of the 3D structures. The developed fabrication process is compatible with the existing design of the photovoltaic retinal implants, and thus immediately translatable into clinical testing.

**Figure 1:**
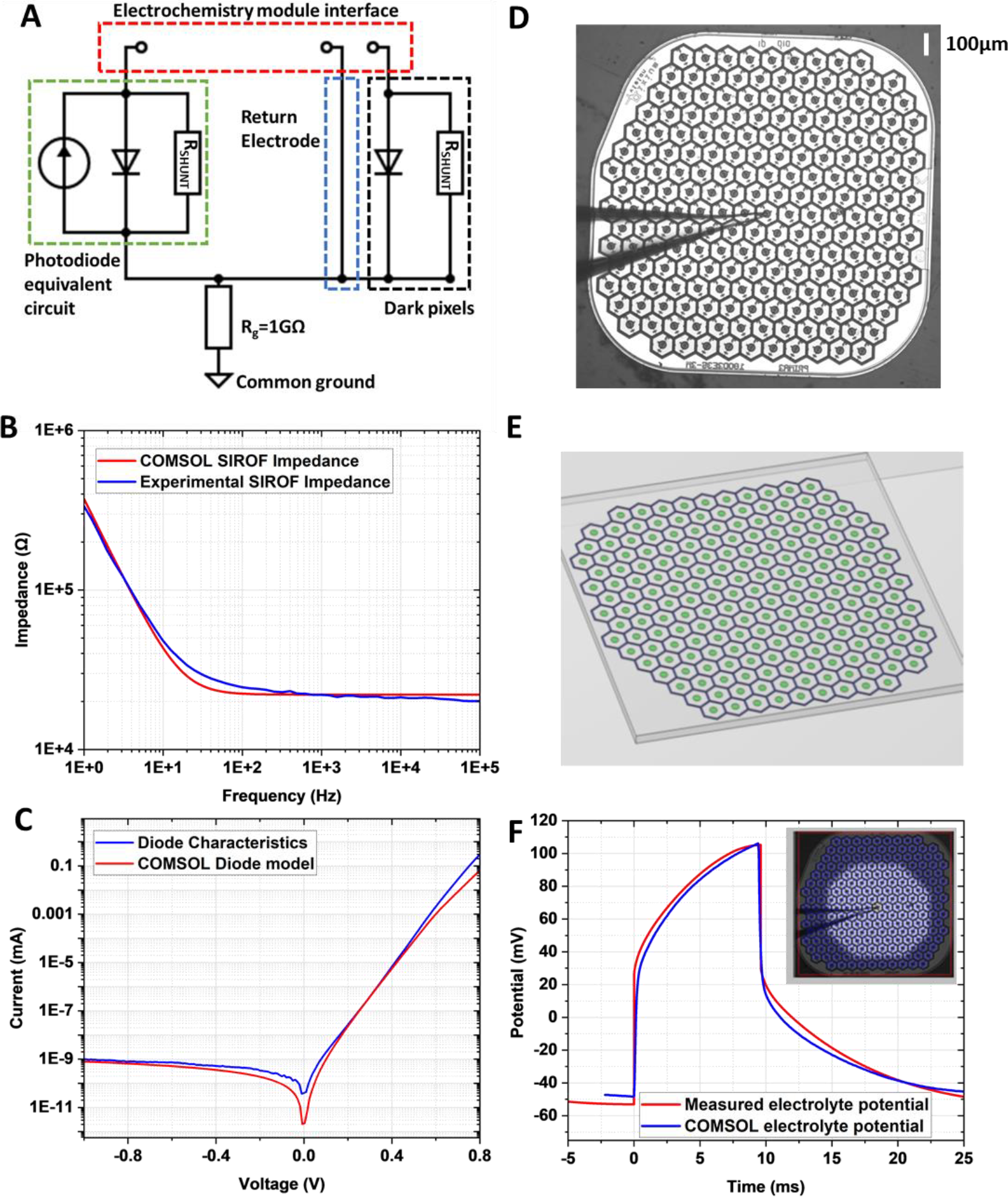
A) SPICE circuit model used to drive the input electrical signals into the COMSOL electrochemistry model. Each modelled pixel is represented by an equivalent circuit model coupled to the stimulation (active) electrode in the electrochemistry solution, where the hexagonal return electrodes are connected through a common terminal. Illuminated pixels also have a current source. B) Modelled and experimental measurements of electrode impedance across frequency for a 100mV pk-to-pk sinusoidal input to an 80 µm diameter active SIROF. C) I-V characteristics of a diode in the SPICE circuit model compared to experimental results from fabricated photodiode array [4]. D) Experimental set up for electrolyte potential measurement. A PRIMA implant was submerged in NaCl solution (1.52 mS/cm), and a micro-pipette electrode used to measure electrolyte potential at 17 µm above the device surface. E) This experimental setup was replicated in the electrochemistry model in COMSOL. F) Experimental and modelled electric potential 17 µm above the implant under spot illumination (diameter = 1000 µm, λ=880nm, irradiance = 3mW/mm^2^, pulse duration 10 ms).

## 2. Methods

### 2.1 Modelling

The finite element analysis tool, COMSOL Multiphysics, was used to calculate the potential throughout the modelled conductive domain. Analysis was carried out using the electrochemistry module in three dimensions to simulate the electrolyte regions and the electrode-electrolyte surface boundaries. Electric potential was computed by coupling the Poisson equation for current density in the electrolyte with the Nernst-Planck equation for flux of charge carriers, assuming electroneutrality and negligible charge carrier gradients [16]. Reactions at electrode surfaces were modelled using the Butler-Volmer equation, describing anodic and cathodic reactions. The electrochemistry module was coupled to a circuit model in COMSOL, which represented individual photodiodes, driving current to active electrodes in illuminated pixels, as well as a path to the interconnected return electrodes. Using these coupled models, allows simulation of the access resistance, double layer capacitance, electrode kinetics and electrolyte potential through the electrochemistry module, whilst the circuit model can drive the stimulation pattern and allow electrode surfaces to have floating potentials, without defining their boundary conditions, which change over time [17]. Using this approach, current is injected into the electrochemical system, as defined by the circuit model, and is collected by either the return electrodes, or the adjacent active (stimulation) electrodes, the potential of which is determined by the circuit dynamics (Fig. 1(a)).

### 2.2 Model Verification

To calibrate the model we compared: 1) modelled electrode impedance values to electrochemical impedance spectroscopy (EIS) measurements from microelectrode structures; 2) the current-voltage characteristics for the modelled photodiode to the experimental results from our photovoltaic device [4] and 3) the computed electric potential to the measured voltage pulses in electrolyte generated by PRIMA implants.

The impedance of a SIROF (sputtered iridium oxide film) electrode surface was modelled using the electrical impedance spectroscopy (EIS) component of the electrochemistry module in COMSOL. Based on our previous measurements, the SIROF capacitance was set to C_Sirof_ = 8.52 mF/cm^2^, for a Sodium Hypochlorite cleaned surface [18]. SIROF is used for the active electrode and return electrodes due to its high charge injection capacity (CIC) compared to other electrode materials [26]. Even though reversible Faradaic reactions contribute to the high capacitance of SIROF, known as the pseudo-capacitance, we combine the double-layer and faradaic capacitances as C_DL_ in COMSOL. Conductivity of the electrolyte domain was set to 2.83 mS/cm to match the conductivity of the diluted phosphate buffered saline solution used in ex-vivo experiments. An exchange current density of 1 mA/cm^2^ was set for the SIROF electrode interface [19]. A frequency sweep was performed and plots of the absolute value of impedance against frequency showed close agreement with the experimental results – Fig. 1(b).

As shown in Fig. 1(a), the pixel equivalent circuit is modelled as a current source, a diode and a shunt resistor in parallel. The I-V characteristics of the diodes used in this equivalent circuit were set to match the photodiodes of the retinal prosthesis detailed in reference [4]: junction capacitance of 30 pF, ideality factor of 1.5, responsivity of 0.51A/W (Fig. 1(c)).

### 2.3 3D Electrode model

With the circuit model and electrode/electrolyte interfaces set, we modelled an array of 100 µm pixels, matching a PRIMA implant – Figures 1(d) and (e), and evaluated the electrolyte potential 17 µm above the device. This was compared to experimental recordings via pipette electrode positioned 17 µm above the PRIMA device in a diluted saline solution (conductivity = 1.52 mS/cm) [20] and illuminated at 3 mW/mm^2^ with pulses of 9.6 ms in duration. As shown in Figure 1(f), the simulated output closely matches the experimental waveform, demonstrating that the model accurately represents the photovoltaic arrays in electrolyte.

To model the 3D honeycomb arrays, 24 µm tall walls of 4 µm width were added on top of the pixel return electrodes with a 22 µm pitch. Each pixel contained a central active electrode, 9 µm in diameter and 400 nm in height. These 3D structures were positioned on a 30 µm thick substrate, which represents the silicon photovoltaic implant, and placed within a 150 µm thick layer (conductivity 1 mS/cm [7]) to represent the retina, within a 1 mm cube representing the vitreous (conductivity 15 mS/cm [27]). A 500 µm inner radius and 510 µm outer radius ring electrode surrounded the modelled array to act as a distant return electrode. The honeycomb walls were modelled as gold, while the active electrodes and caps on top of the walls, modelled as 400 nm thick SIROF. Current pulses are defined in the circuit model, which then determines the current and voltage on active and return electrode interfaces in the electrochemisty module. All other surfaces are defined as electrically insulating (Neumann boundary conditions). Due to the shunt resistors and the diode conductivity under suffcient bias, the active electrodes in non-illuminated pixels (both honeycomb and pillar models) can collect current just like the return electrode mesh in the honeycomb model. Pillar active electrodes were modelled by placing the 9 µm diameter SIROF active electrode on top of a 35 µm high Au pillar (same diameter) and using the 0.5 mm radius ring as a common return electrode.

The magnitude of the current source in each pixel was calculated based on a responsivity of 0.5 A/W [4], the photoactive area of a pixel and an illumination of 1 mW/mm^2^ (at µ=880 nm). A shunt resistor is included in each pixel to help discharge the pixel between the light pulses (30 Hz, 4 ms pulse width). The optimal value of the shunt resistor depends on the pixel size. Using a value of approximately 5 times the access resistance, a shunt of 720 kΩ was selected for 100 µm pixels. When modelling the 20 µm pixel arrays, a shunt value of 4 MΩ was selected using the same criteria. The side walls (C_DL_=14-100 µF/cm^2^) and SIROF caps of the return electrodes in each pixel are connected to the terminals of the circuit model, and all the return electrodes are connected together in one common mesh. All current applied through the stimulation (active) electrode is collected by the other electrode surfaces, such that the total charge in the system is conserved.

### 2.4 Fabrication of 3D electrodes

We have previously described the fabrication process for planar photovoltaic retinal implants [4]. Here we detail fabrication processes and procedures for integration of the 3D electrode structures, building upon established fabrication procedures of the planar devices. These electrodes are electroplated onto the photovoltaic arrays after the fabrication of photodiodes, but before the electrode interface material (SIROF) is deposited.

In order to develop this process on a protoype wafer, we patterned the active and return electrode structures in a Ti:Au layer (50 nm:200 nm) on blank 4-inch silicon wafers (p-doped) using a lift-off process (500 nm layer of LOR-10B, followed by a layer of Shipley 1805 photoresist). These active and return electrode structures, used as starting points for electroplating the 3D devices, were interconnected across the entire wafer, allowing simultaneous electroplating. Dimensions of the electroplated structures were chosen to match those used in photovoltaic subretinal prostheses, where each hexagonal pixel consisted of a disc electrode in the middle and a circumferential electrode on the edge [4]. Each array was 1.5 mm in diameter, comprised of pixels of 55, 40, 30 or 22 µm in width. For honeycombs, the circumferential electrode of each pixel was electroplated into vertical walls of widths 5.5 µm (55 µm pixels), 4.5 µm (40 µm pixels) and 4 µm (30 and 22 µm pixels). For the pillar design, the disk electrode in every pixel was electroplated into a pillar, with diameters of 22, 16.5, 11.5 and 8.5 µm for pixels of 55, 40, 30 or 22 µm, respectively.

A thick high-aspect ratio negative photoresist (KMPR-1025) was used to define the mask for electroplating honeycombs or pillars. The electroplating pattern was transferred using a contact aligner (Karl Suss MA6), and development was carried out with a TMAH-based developer. Patterned wafers were fixed into a custom-made, PTFE wafer holder, which isolated the back surface and edges of the wafer, so that only the desired areas were exposed to electroplating solution (NB Semiplate AU 100^TH^, NB Technologies, Bremen, Germany). A hollow handle provided electrical contact to the Ti:Au layer on the wafer surface, while a platinized titanium mesh, positioned parallel to the wafer surface, was used as the anode. A hot plate kept the solution at a temperature of 30°C and a stirrer provided constant agitation at 40 rpm. A constant current density of 1 mA/cm^2^ was applied, providing a plating rate of 3 µm per hour. After electroplating up to the desired height, the solution was removed, and the wafer rinsed with DI water. The KMPR-1025 electroplating template was removed using PG remover at 80 °C, leaving the desired 3D honeycomb or pillar pattern in gold.

To coat the tops of the walls and pillars with a SIROF layer, a lift off process was used. Once electroplated, the wafers were spray-coated in photoresist (50 µm thick) and processed through a repetitive cycle of underexposure and development to remove the resist, in a layer by layer fashion, until the top of the electroplated metal structures were revealed. The top surface of the electroplated structures was then sputter-coated with Ti:SIROF (40 nm:436 nm), providing a high-capacitance material for the electro-neural interface. The fabrication procedure concludes by dissolution of the remaining photoresist, revealing the 3D walls and pillars with SIROF on the top surface and exposed Au on side walls. Backside grinding is then carried out to thin each wafer to 30 µm. Soaking in acetone released each individual 3D array from the supporting grinding tape.

Honeycomb walls were fabricated to 24 µm in height (the approximate thickness of the inner nuclear layer), while pillar height was set to 35 µm, to match the debris layer thickness in AMD patients [15].

### 2.5 Animals, surgical procedures and tissue processing

All experimental procedures were approved by the Stanford Administrative Panel on Laboratory Animal Care (APLAC) and conducted in accordance with the institutional guidelines and conformed to the Statement for the Use of Animals in Ophthalmic and Vision research of the Association for Research in Vision and Ophthalmology (ARVO). Royal College of Surgeons (RCS) rats were used as a model of photoreceptor degeneration. Total of N = 3 animals were implanted with pillar arrays after the age of P180 to ensure complete degeneration of the photoreceptors. As previously described [3], animals were anesthetized with a mixture of ketamine (75 mg/kg) and xylazine (5 mg/kg) injected subcutaneously. A 1.5 mm incision was made through the sclera and choroid 1 mm posterior to the limbus. The retina was detached with an injection of saline solution, and the implant was inserted into the subretinal space at least 3 mm away from the incision site. The conjunctiva was sutured with nylon 10-0, and topical antibiotic (bacitracin/polymyxin B) was applied on the eye postoperatively. The eyes were collected 8 days later and fixed in 4% PFA. The retinal whole mount was stained with DAPI nuclear marker, imaged by LSM 880 confocal microscope (Zeiss LSM 880, Germany) and reconstructed using ImageJ (Fiji) and MATLAB 2021b (Mathworks, Inc., Natick, MA).

## 3. Results

### 3.1 Modelling the neural stimulation

After validating the electrochemical model by comparison with experimental results, as described in sections 2.2 and 2.3, we investigated the effect of three-dimensional structures on the electric field generated by 20 µm pixels. An array of 59 pixels, 20 µm in pitch were modelled as described in section 2.3, and simulations carried out using planar, honeycomb and pillar geometries. A 4 ms stimulation pulse was applied to the current source in the circuit model (Figure 1(a)), with an amplitude calculated for illumination of 1 mW/mm^2^, responsivity of 0.51 A/W [4] and the photoactive area of a 22 µm photovoltaic pixel – 217 µm^2^. There is a range of double layer capacitance values for gold in the literature, depending on the surface smoothness and preparation. For our modelling, we have selected a C_DL_ = 56 µF/cm^2^ from [21], but also explored the effect of lower (14 µF/cm^2^) and higher (100 µF/cm^2^) values of C_DL_. An exchange current density of 2 nA/cm^2^ was used for the gold in electroplated walls and pillars [23].

Figure 2(a) depicts the electric potential in electrolyte in front of the planar and honeycomb arrays at the end of a 4 ms pulse. Diagram of a bipolar cell (BC) in front of the array (and migrated into the well) is shown to scale in this cross-section. Axonal terminals of BCs are in the middle of the inner plexiform layer (IPL), up to 65 µm above the surface of the array, while cell somas reside inside the honeycombs [7]. The electric potential calculated through the modelled retinal volume is plotted with respect to the middle of the IPL. Previous work has shown that for a 4 ms anodic pulse, a potential difference of at least 4.3 mV across the cell body from soma to axonal terminals is required to generate a retinal response [10]. This stimulation threshold is indicated in figure 2(a) as the cyan contour, demonstrating the field enhancement effect produced by the 3D structure. Within the honeycomb walls, this region extends towards the top of the wall, compared to a flat planar array, where it is confined to much smaller volume above the active electrode. Even though the three-dimensional return walls in our simulation are modelled as an electrically conducting surface, their small capacitance (14-100 µF/cm^2^) results in this being a relatively high impedance path, compared to the SIROF cap electrodes (>4 mF/cm^2^). Figure 2(b) shows this effect in a model, where an initial spike of current flows into the sidewall at the beginning of a pulse, but then rapidly decreases as the sidewall capacitance charges up. Within 0.3 ms, majority of current starts flowing through the much higher capacitance of SIROF cap electrode, providing the vertical current alignment, matching the orientation of bipolar cells. Notably, the cathodic potential on the Au sidewalls is approximately −40 mV during stimulation, well below the electrochemical reactions’ threshold for Au [21].

**Figure 2:**
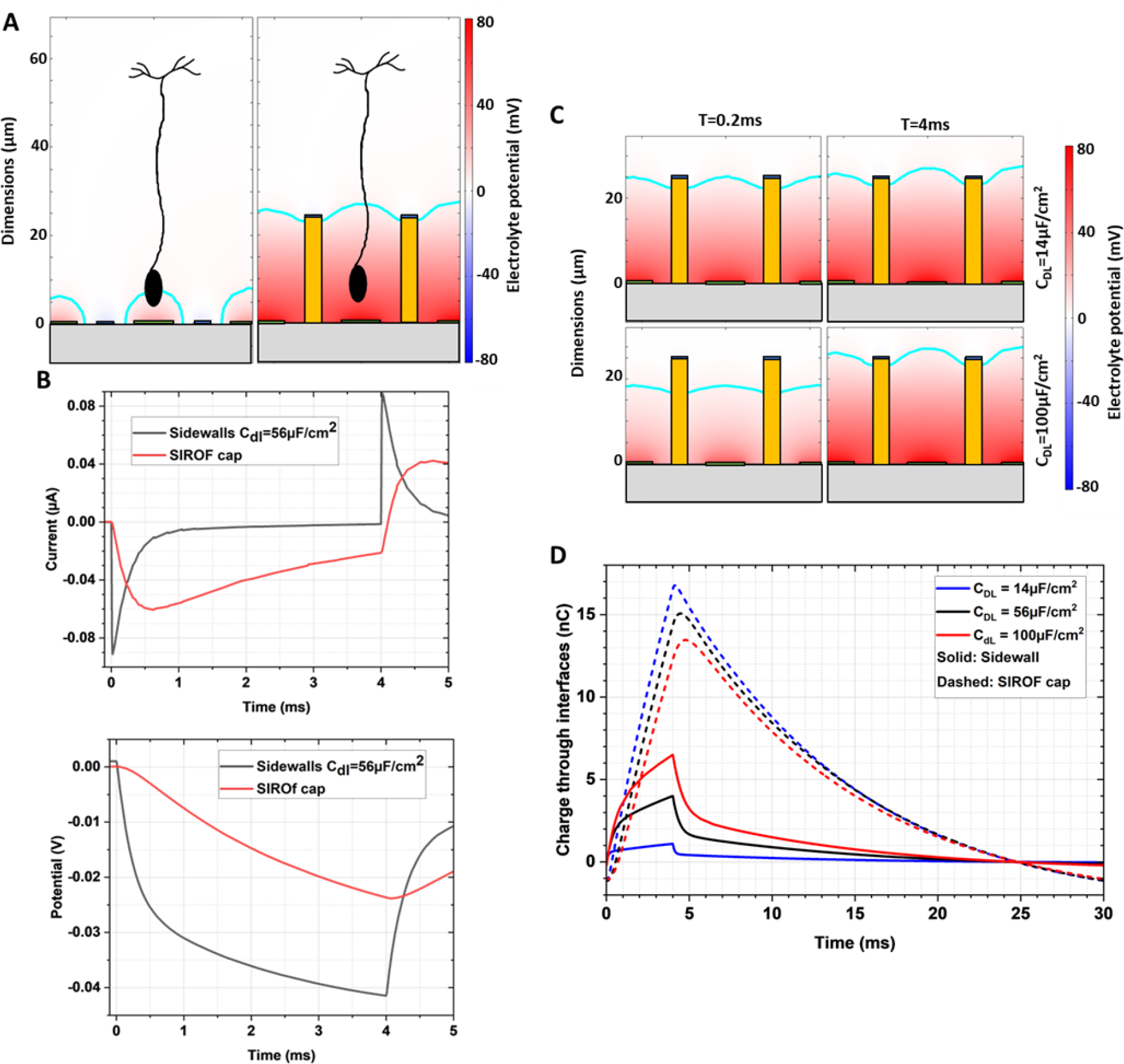
A) Left: Electric field penetration is limited for planar devices with small pixels, due to the proximity of the active and return electrodes. Right: Placing the return electrodes on top of vertical walls, helps extend the field vertically and permits stimulation with small pixels. Electrolyte potential is depicted with respect to the middle of the IPL, where bipolar cell axons terminate. The cyan contour indicates the region above an assumed stimulation threshold of 4.3mV. B) Left: COMSOL model of the current flow across the return electrode structure. The electroplated sidewalls are modelled as electrical conductors meaning that current can be sinked through this interface. Initially, current flows into the sidewalls, but then the high capacitance SIROF coating that caps the return structure, becomes the preferred current path. Right: The potential drop across the gold sidewall interface does not reach the levels where the onset of oxygen reduction reactions can occur [21]. C) Different electrode materials exhibit a range of capacitances which affects current flow into the sidewall. Two examples are shown for Au (14 µF/cm^2^) and Pt (100 µF/cm^2^) interfaces. Across this range of surface capacitances, current flow into the SIROF cap still dominates with little difference in the electrolyte potential profile at the end of a 4ms pulse. D) The total charge collected by the sidewalls and SIROF cap. With increasing sidewall capacitance, the amount of charge collected by sidewalls (over a 4ms pulse) increases, with a corresponding decrease in the charge collected through the SIROF cap.

As mentioned above, there is a range of C_DL_ values for gold in literature, and we investigated the effect of changing C_DL_ from 14 to 100 µF/cm^2^. Figure 2(c) shows electric potential with honeycombs at the onset and end of a 4ms pulse. With a double layer capacitance set to 100 µF/cm^2^ (bottom plots), the field penetration depth is initially restricted to ~10-15 µm. However, by the end of the pulse, current flows once again predominately through the SIROF cap. Figure 2(d) shows the time course of this process in terms of the charge collected by different surfaces. As the sidewall capacitance increases, the proportion of total charge collected by the sidewall also increases, from 6% for C_DL_ = 14µF/cm^2^, 27% for C_DL_ = 56µF/cm^2^, to nearly 50% for C_DL_ = 100µF/cm^2^.

The model also demonstrates the effect of charge redistribution across the return electrode surface. When 19 pixels are illuminated, as shown by dashed circle in Figure 3, the current density on the top surface (SIROF) is initially confined to the return electrode surface within illuminated area. From 1 ms onwards, current begins to redistribute more evenly, recruiting the return electrode surfaces of non-illuminated pixels. This also occurs with the current returning in opposite polarity after the light pulse is over (t=5 and 10 ms). This dynamic is summarised in a plot for electrodes at the centre pixel (1), at the edge of illuminated circle (2) and in the non-illuminated area (3).

**Figure 3:**
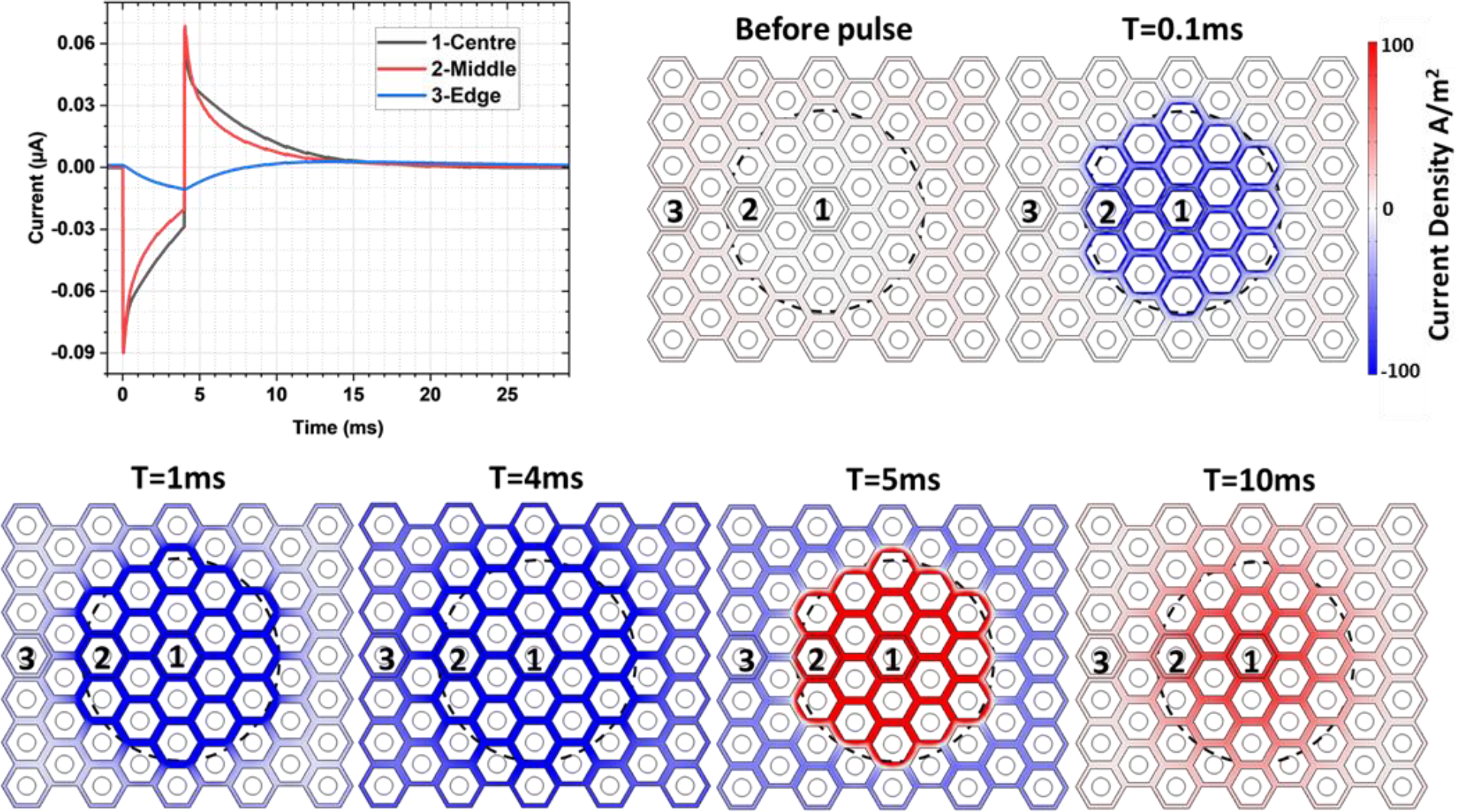
When a spot (indicated by the dashed circle) is illuminated by a 4ms pulse, current is initially collected by the adjacent return electrodes, but over time it is redistributed across the entire return electrode mesh. Similar redistribution occurs after the light pulse, with the return interfaces trending back towards equilibrium. Current density on active electrodes is not shown here.

With 35 µm tall, electroplated Au pillars acting as active electrodes, adjacent non-illuminated active electrodes can act as returns, collecting the injected current through the shunt resistor or via diodes under sufficient bias. With pillars, field is not shaped vertically as in honeycombs, but rather exhibits a spherical expansion, similar to the disc electrodes. Figure 4(a) shows the debris layer under the INL, which pillars are expected to penetrate, and evolution of the threshold potential difference (4.3 mV) during the 4 ms pulse modelled using both gold and platinum (C_DL_=100 µF/cm^2^) pillars. After initial charging of the side walls, the threshold contour becomes (and stays for the remainder of the pulse) localized to the top of the pillar, surrounding the SIROF cap. The current and voltage pulses are shown in Figures 4(b) and (c). Notably, the pillar sidewall potential does not exceed 100 mV, below the level for electrochemical reactions, such as oxygen reduction onset and hydrogen peroxide evolution (~100 mV). Figure 4(d) shows the percentage of the INL volume above the stimulation threshold (4.3 mV [7]) during a 4 ms pulse. Previous work has shown that a visual response can be evoked when ~8 % of the targeted volume is above the stimulation threshold [7].

**Figure 4:**
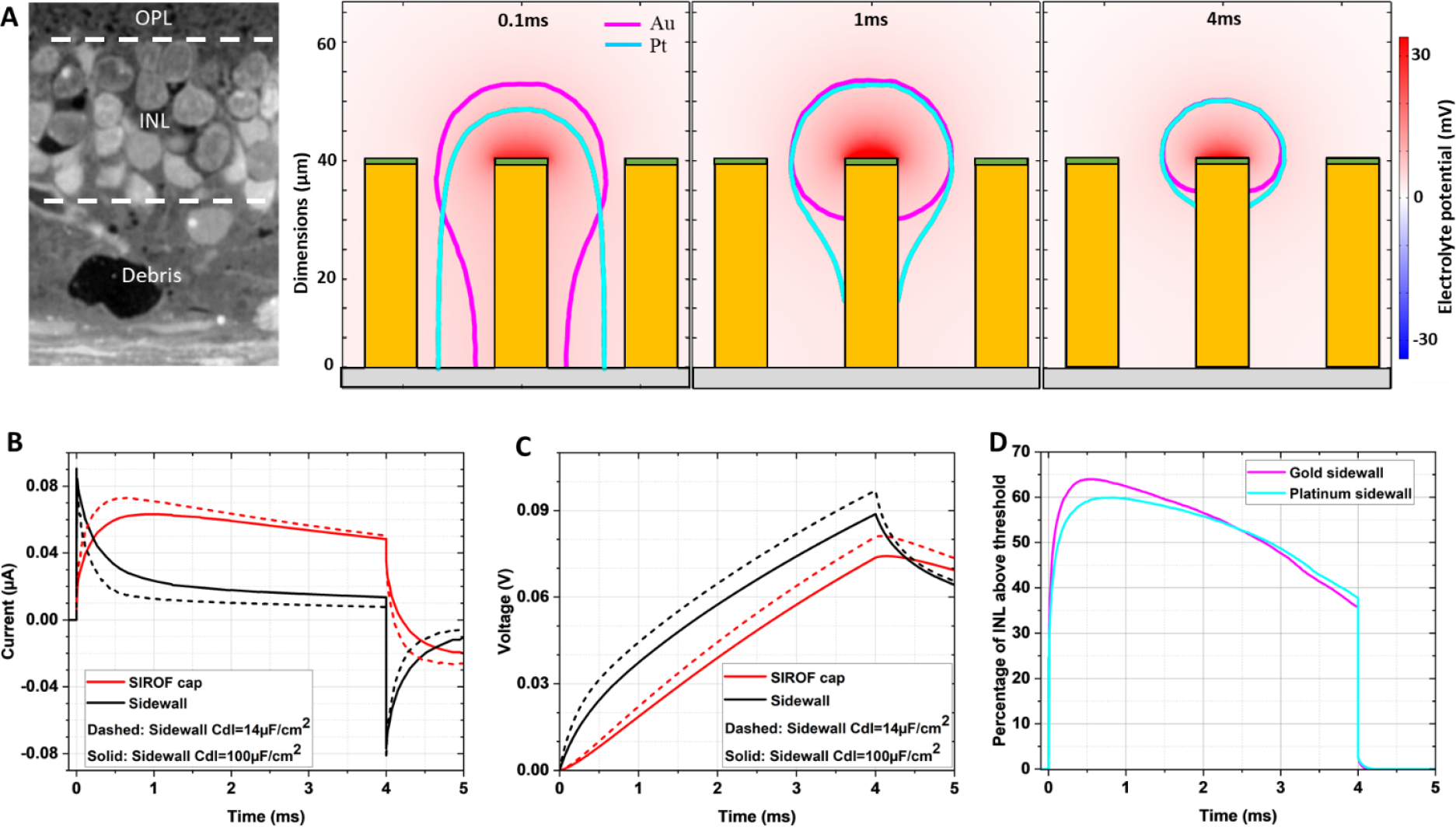
A) The degenerated retina can exhibit debris layers up to 40 µm thick in human patients. Electroplated pillars can penetrate this layer to deliver current to the targeted bipolar cells. The colour map indicates the electrolyte potential with respect to the middle of the IPL for a gold pillar and the purple contour indicates the volume above a stimulation threshold of 4.3 mV. Cyan contour indicates the same volume for a platinum pillar. B) Stimulation through the pillar electrode, initially results in current flow out of the sidewall. However, this interface quickly saturates, and the preferred current path is then via the SIROF cap, where the majority of current is injected. C) Potentials on the sidewall and SIROF cap do not exceed safety thresholds of +100 mV. D) Percentage of INL volume above the stimulation threshold of 4.3 mV during a 4 ms pulse. Previous work [11] has seen that VEP threshold corresponds to 8.3% of the targeted INL volume above the stimulation threshold.

Comparison between Pt and Au pillars demonstrates the effect of sidewall capacitance, altering the potential in electrolyte, especially at the beginning of a 4 ms pulse – figure 4(a). This results in a larger current through the Pt sidewall - figure 4(b) and a lower potential on the Pt sidewall - figure 4(c). In consequence, a greater fraction of current passes through the pillar sidewall and slightly lower fraction of the targeted volume is above the stimulation threshold – figure 4(d).

### 3.2 Fabrication

Our electroplated honeycombs and pillars are shown in Figure 6. A cross section of the photoresist pattern used as an electroplating mask is shown in Figure 5(a). The 24 µm tall electroplated wall structures, integrated with an underlying device pattern to alignment accuracy of 1 µm, are shown in Figures 5(b-d). Electroplated pillars of 35 µm in height with diameters matching the size of the active electrodes in the existing photovoltaic arrays [4] are shown in Figures 5(e-h).

**Figure 5:**
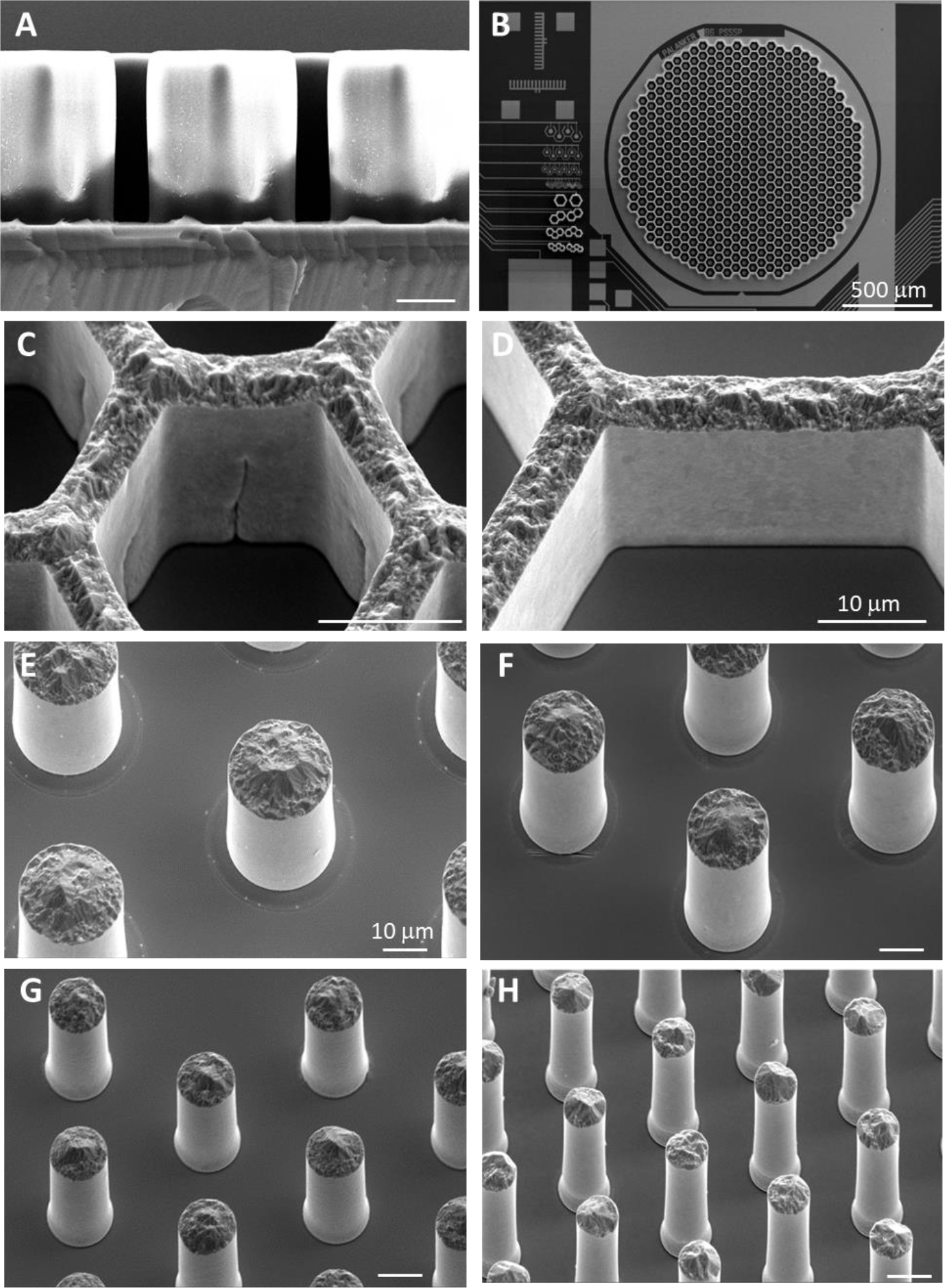
A) A high-aspect ratio negative photoresist pattern is used to define the electroplated three-dimensional walls. B) After the wafer scale electroplating process, the photoresist is stripped, leaving 25-35µm high electroplated structures. This process has been adapted to integrate with the photovoltaic retinal prosthesis. C) and D) Fabricated structures with smooth sidewalls, up to 24µm in height and widths of 4µm. These dimensions are tall enough to integrate with the targeted cells and narrow enough to limit the shadowing of the underlying photoactive area. E), F), G) and H) the fabrication process was adapted to produce pillar electrodes, capable of penetrating past a retinal debris layers, with heights up to 35µm and widths of 23µm, 17µm, 13µm and 8.5µm.

**Figure 6:**
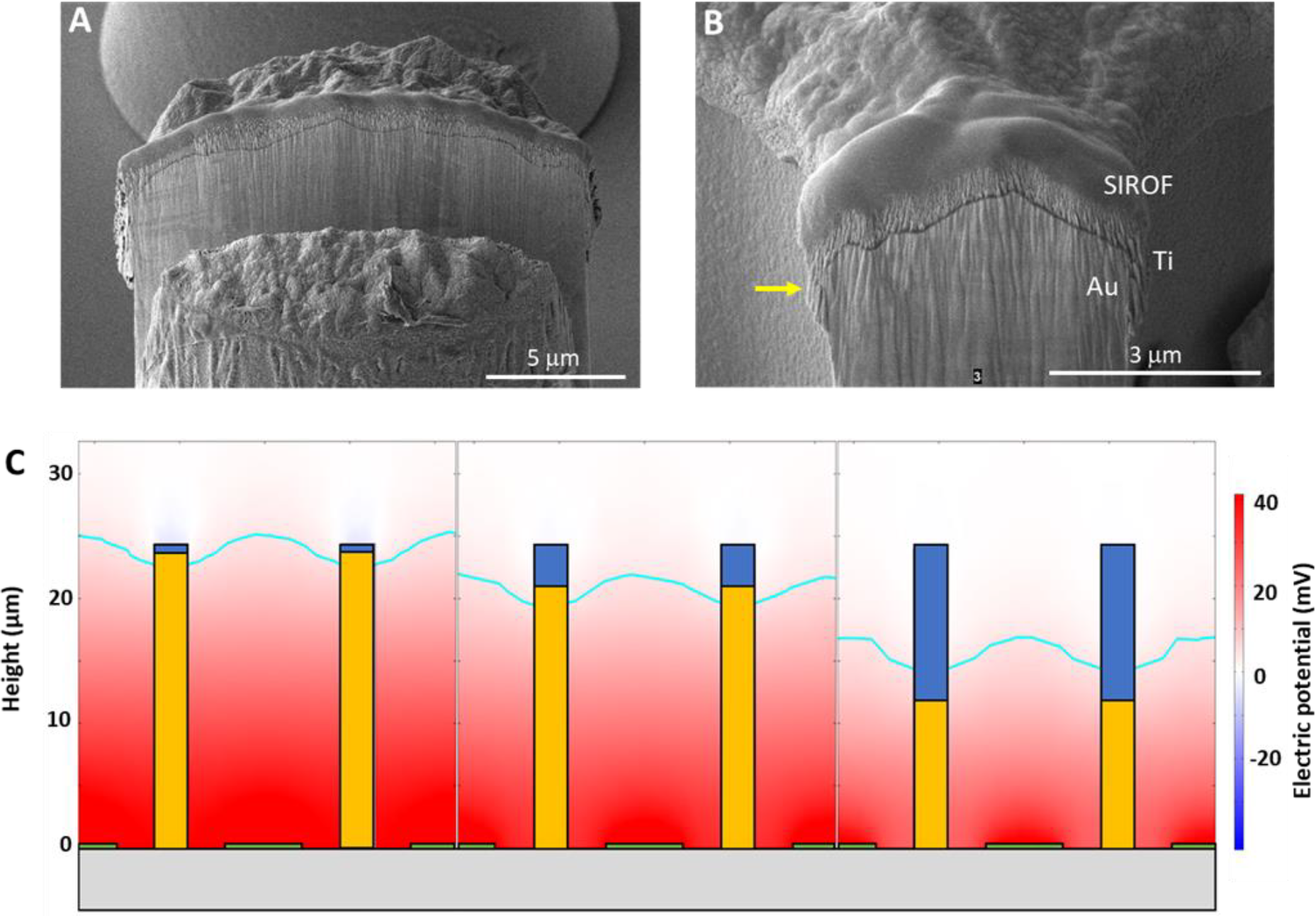
Focused Ion Beam milling of the SIROF coating on top of electroplated pillars (A) and honeycomb electrodes (B). C) Deposition of SIROF on top of electroplated gold structures can result in an overhang of a few µm (pointed by an arrow), due to non-uniformities in the lift-off process. Models indicate that an overhang of 4µm results in a reduction of 7% in the targeted cell volume above the stimulation threshold, whilst an overhang of 12µm would result in a reduction of 17% of this volume.

The top surface of walls and pillars were then coated with a 400nm thick SIROF layer, shown in Figure 6(a). Due to variations in height (±0.8 µm) of electroplated structures across a 4-inch wafer and variations in thickness of the photoresist used for lift-off, a SIROF overhang of 1-4 µm in height was observed (pointed by the yellow arrow in Figure 6(b)). We analysed the effect of such an overhang on electric field within the honeycombs. As shown in Figure 6(c), an overhang of 4 µm decreases the field penetration depth by 7%, compared to the ideal case with a 400 nm SIROF cap. This trend continues, as shown for an extreme overhang covering half of the wall height. Such a large SIROF overhang was not seen in our fabrication results.

We assessed the mechanical stability of these 3D devices by testing their release from the carrier wafer and a subretinal implantation in rats. Since the connected honeycomb walls are much stronger than individual pillars, we focused our effort on evaluating the mechanical stability with pillars. As shown in Figure 7(a), all the pillars on 30 µm pixels still stand after the implants release from the carrier wafer. Figures 7(b) and (c) show a cross-section of a confocal image of the device in subretinal space after implantation and explantation 8 days later. Pillars are still standing, and migration of the inner retinal cell into the voids of the array in Figure 7(b) indicates feasibility of the tight integration of these 3D structures with the retinal tissue for close proximity to the target neurons. Further details of the retinal integration with 3D implants can be found in [24].

**Figure 7:**
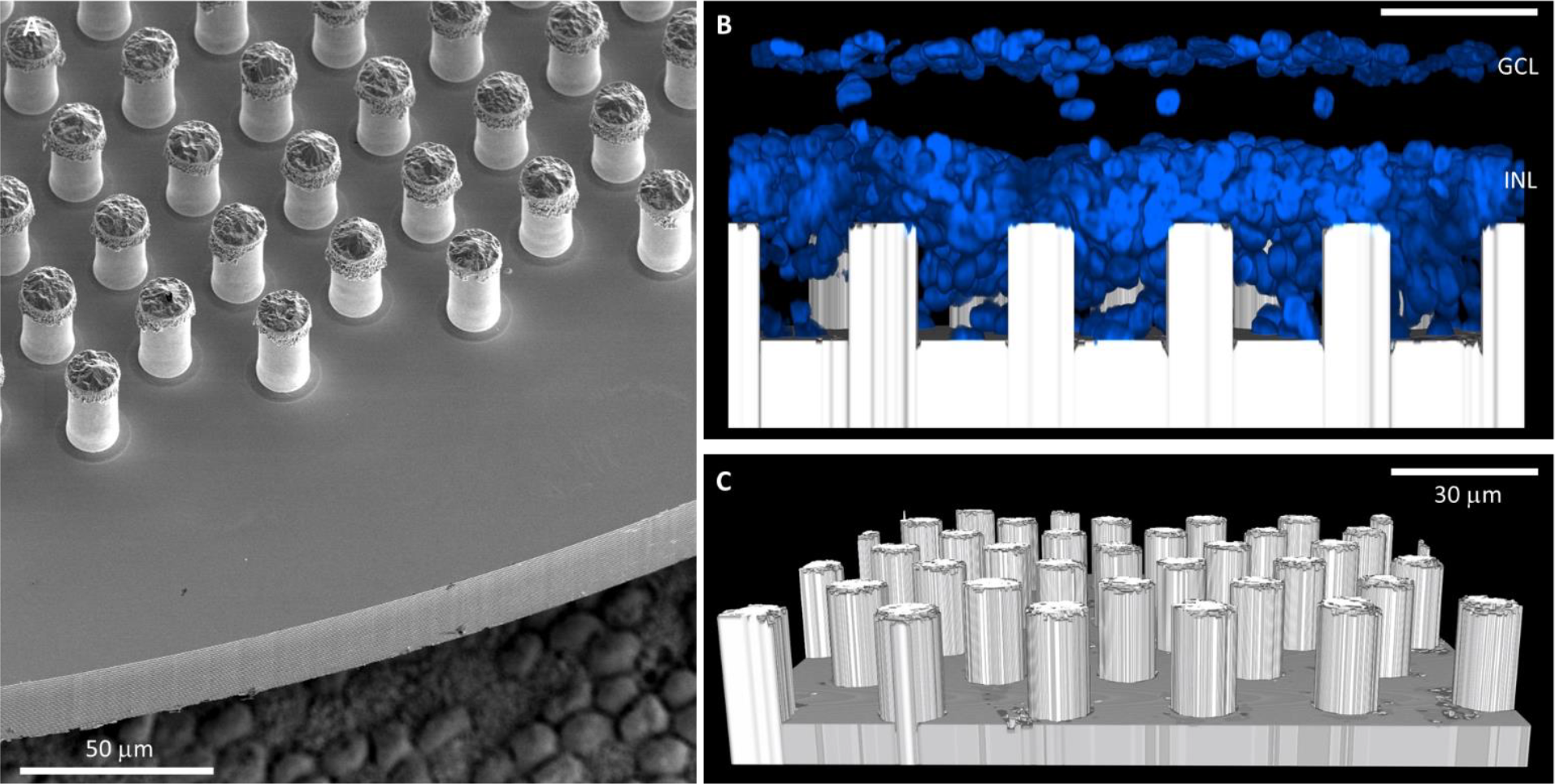
A) Released array with 30 µm high pillar electrodes, capped with a SIROF (array is placed on top of retinal pigment epithelium cells to give an idea of scale). B): Cross section of the explanted pillar structures on 30 µm pixels, integrated with the rat retina. DAPI indicates cell bodies in blue. 30 µm high pillars survived release of individual arrays, implantation and explantation.

## 4. Discussion

3D electrode structures are essential for high resolution subretinal implants since they improve proximity and enable more focused stimulation of the second-order neurons in the retina. However, fabrication of such structures on semiconductor devices, which rely on planar manufacturing processes, is challenging. Here, we have shown how these devices can be produced using conventional lithographic techniques coupled to a gold electroplating process that enables high-aspect ratio structures in the form of either honeycomb walls or pillars. The process is compatible with the post-processing of wafer-level devices and is robust enough to withstand the mechanical thinning of the wafers, release of the individual devices and implantation in rats. Furthermore, the 3D electrode structures integrate well with the retinal tissue, with histological results showing migration of the inner retinal neurons into the spaces between electrodes, promoting close proximity between stimulation electrodes and cells.

An important consequence of this fabrication process is the exposed metallic sidewalls of the 3D structures. Ideally these would be insulated with a dielectric thin film, however, this is complicated with high-aspect ratio 3D structures and the non-conformal coating of most deposition techniques. Modelling indicates that smooth (low capacitance) Au sidewalls do not provide significant charge transfer, which is dominated by the (high capacitance) SIROF electrodes on the top surface of the 3D structures. The Au sidewalls are charged predominantly during a submillisecond time window on the rising and falling edges of the stimulation pulse. The magnitude depends upon the double layer capacitance, with 5-25% charge loss over the likely range of C_DL_ values (14-56 µF/cm^2^). On pillar active electrodes, the electrode potentials required to deliver these stimulation pulses are positive and quite low (<100 mV). On honeycomb return electrodes, these potentials are negative, and even lower (about −40 mV), well below the thresholds for oxygen reduction (−300 mV), H_2_O_2_ evolution (−300 mV) and H_2_ evolution (−700 mV) [21]. The models indicate that as long as the potential at the Au interface is kept away from the redox reaction levels, then the low capacitance of these exposed side walls obviates the need for their electrical insulation, greatly simplifying the microfabrication process.

Althought the models indicate that the surface potentials and charge densities on the Au surfaces are below the commonly quoted values for detrimental reactions in vivo, a more common electrode material is platinum. Pt electrodes are a clinical standard in very successful long-term neural implants, such as deep brain stimulators and cochlear implants [25]. This makes Pt an attractive material for 3D electrodes, however, there is a caveat. Higher C_DL_ of Pt (100 µC/cm^2^) compared to Au means more charge is driven across the exposed walls during stimulation, reducing the current delivered via the SIROF cap to the target cells. However, our models indicate that more than half of the targeted cells can still be safely driven above the stimulation threshold (4.3 mV) using modest irradiance values (~1mW/mm^2^), when the potential on sidewalls does not exceed the 100mV threshold for the onset of oxygen reduction [21]). However, if stronger stimulation is applied (e.g. 3 mW/mm^2^ used with the PRIMA implants in clinics), it may result in higher voltages, exceeding the threshold of irreversible electrochemical reactions. To prevent them, atomic layer deposition (ALD) coatings could be introduced. Such insulating coatings on the sidewalls could eliminate the possibilty of any reactions and further concentrate the electrical current throught the SIROF caps.

These 3D electrode structures electroplated on top of the planar subretinal arrays hold promise for either shaping the electric field vertically (honeycombs) or raising the stimulating electrode to the target neuronal layer (pillars), both of which improve the efficiency of retinal stimulation and will help facilitate high-density neuromodulation.

## Acknowledgements

Studies were supported by the National Institutes of Health (Grants R01-EY-027786 and P30-EY-026877), the Department of Defense (Grant W81XWH-2210933), AFOSR (Grant FA9550-19-1-0402), Wu Tsai Institute of Neurosciences at Stanford, and unrestricted grant from Research to Prevent Blindness. Photovoltaic arrays were fabricated at the Stanford Nano Shared Facilities (SNSF) and Stanford Nanofabrication Facility (SNF), which are supported by the National Science Foundation award ECCS1542152.

K.M. was supported by a Royal Academy of Engineering Chair in Emerging Technology, UK.

The research was funded by the Royal Academy of Engineering, the Rhona Reid Foundation and the EPSRC.

## References

[1] Schwartz D, et al 2014 Principles of Tissue Engineering (Fourth Edition) Pages 1427–1440 ISBN 9780123983589

[2] Smith W, Assink J, Klein R, Mitchell P and Klaver C 2001 Risk factors for age-related macular degeneration: pooled findings from three continents Ophthalmology 108 697–704

[3] Lorach H et al 2015 Photovoltaic restoration of sight with high visual acuity Nat. Med. 21 476–82

[4] Vertical-junction Photodiodes for Smaller Pixels in Retinal Prostheses. T.W Huang, T.I Kamins, Z.C. Chen, B. Wang, M. Bhuckory, L. Galambos, E. Ho, T. Ling, S. Afshar, A. Shin, V. Zuckerman, J.S Harris, K. Mathieson, D. Palanker. J. Neural Eng. 18 036015 (2021).

[5] Palanker D, Vankov A, Huie P and Baccus S 2005 Design of a high-resolution optoelectronic retinal prosthesis J. Neural Eng. 2 S105–20

[6] Zhou D D, Dorn J D and Greenberg R J 2013 The Argus II retinal prosthesis system: an overview IEEE Int. Conf. on Multimedia and Expo Workshops (IEEE) pp 1–6

[7] Flores T, Huang T, Bhuckory M, et al. 2019 Honeycomb-shaped electro-neural interface enables cellular-scale pixels in subretinal prosthesis Sci Rep 9 10657 10.1038/s41598-019-47082-y

[8] B. Y Wang et al 2022 J. Neural Eng. 19 055003

[9] Muqit MMK, Mer YL, Holz FG, Sahel JA. Long-term observations of macular thickness after subretinal implantation of a photovoltaic prosthesis in patients with atrophic age-related macular degeneration. J Neural Eng. 2022 Oct 14;19(5):10.1088/1741-2552/ac9645. doi: 10.1088/1741-2552/ac9645. PMID: 36174540; PMCID: PMC9684097.

[10] Flores, T. et al. 2018 Optimization of pillar electrodes in subretinal prosthesis for enhanced proximity to target neurons. J. Neural Eng. 15 036011

[11] Flores T, Huang T, Bhuckory M, et al. 2019 Honeycomb-shaped electro-neural interface enables cellular-scale pixels in subretinal prosthesis Sci Rep 9 10657 10.1038/s41598-019-47082-y

[12] Palanker D V, Huie P, Vankov A, Aramant R, Seiler M, Fishman H, Marmor M and Blumenkranz M 2004 Migration of retinal cells through a perforated membrane:implications for a high-resolution prosthesis Investigative Ophthalmol. Vis. Sci. 45 3266–70

[13] Butterwick A, Huie P, Jones B W, Marc R E, Marmor M and Palanker D V 2009 Effect of shape and coating of a subretinel prosthesis on its integration with the retina Exp. Eye Res. 88 22–9

[14] Ho A C et al. 2019 Characteristics of prosthetic vision in rats with subretinal flat and pillar electrode arrays J. Neural Eng. 16(6) 066027 doi:10.1088/1741-2552/ab34b3

[15] Palanker D, et al 2020 Photovoltaic restoration of Central Vision in Atrophic Age-Related Macular Degeneration Opthalmalogy 27 8 1097–1104

[16] COMSOL Electrochemisty Module, P.34 https://doc.comsol.com/5.6/doc/com.comsol.help.echem/ElectrochemistryModuleUsersGuide.pdf

[17] Chen Z, et al. 2020 Current Distribution on Capacitive Electrode-Electrolyte Interfaces Physical Review Applied 13 01004

[18] Cogan S, Plante T and Ehrlich J 2004 Sputtered iridium oxide films (SIROFs) for low-impedance neural stimulation and recording electrodes Engineering in Medicine and Biology Society IENBS’04 4153-6

[19] Boodts, J.C.F., Trasatti, S. Hydrogen evolution on iridium oxide cathodes. J Appl Electrochem 19, 255–262 (1989).

[20] Chen ZC, Wang BY, Kochnev Goldstein A, Butt E, Mathieson K, Palanker D. Photovoltaic implant simulator reveals resolution limits in subretinal prosthesis. J Neural Eng. 2022 Sep 27;19(5). doi: 10.1088/1741-2552/ac8ed8. PMID: 36055219.

[21] Jiří Ehlich, et al 2022 Direct measurement of oxygen reduction reactions at neurostimulation electrodes, J. Neural Eng. 19 036045

[22] K. Najafi and K. D. Wise, “An implantable multielectrode array with on-chip signal processing,” in IEEE Journal of Solid-State Circuits, vol. 21, no. 6, pp. 1035–1044, Dec. 1986, doi: 10.1109/JSSC.1986.1052646.

[23] Mohajeet B. Bhuckory, Zhijie Chen, Bing-Yi Wang, Andrew Shin, Tiffany Huang, Ludwig Galambos, Efstathios Vounotrypidis, Keith Mathieson, Theodore Kamins, Daniel Palanker Cellular integration with a subretinal honeycomb-shaped prosthesis. (December 16, 2022)

[24] B.Y Wang, Z.C. Chen, M. Bhuckory, T.W. Huang, A. Shin, V. Zuckerman, E. Ho, E. Rosenfeld, L. Galambos, T. Kamins, K. Mathieson, D. Palanker. Electronic Photoreceptors Enable Prosthetic Vision with Acuity Matching the Natural Resolution in Rats. Nature Commun. 13: 6627 (2022).

[25] Cogan SF. Neural stimulation and recording electrodes. Annu Rev Biomed Eng. 2008;10:275–309. doi: 10.1146/annurev.bioeng.10.061807.160518. PMID: 18429704.

[26] A. N. Selner, Z. Derafshi, B. E. Kunzer, et al., “Three-dimensional model of electroretinogram field potentials in the rat eye,” IEEE Transactions on Biomedical Engineering 65(12), 2781–2789 (2018).

